# The Dynamics of Synthesis and Localization of Jumbo Phage RNA Polymerases inside Infected Cells

**DOI:** 10.1101/2023.09.14.557509

**Authors:** Daria Antonova, Viktoriia V. Belousova, Erik Zhivkoplyas, Mariia Sobinina, Tatyana Artamonova, Innokentii E. Vishnyakov, Inna Kurdyumova, Anatoly Arseniev, Natalia Morozova, Konstantin Severinov, Mikhail Khodorkovskii, Maria V. Yakunina

## Abstract

A nucleus-like structure composed of phage-encoded proteins and containing replicating viral DNA is formed in *Pseudomonas aeruginosa* cells infected by jumbo bacteriophage phiKZ. The PhiKZ genes are transcribed independently from host RNA polymerase (RNAP) by two RNAPs encoded by the phage. The virion RNAP (vRNAP) transcribes early viral genes and must be injected into the cell with phage DNA. The non-virion RNAP (nvRNAP) is composed of early genes products and transcribes late viral genes. In this work, the dynamics of phage RNAPs localization during phage phiKZ infection was studied. We provide direct evidence of PhiKZ vRNAP injection in infected cells and show that it is excluded from the phage nucleus. The nvRNAP is synthesized shortly after the onset of infection and localizes in the nucleus. We propose that spatial separation of two phage RNAPs allows coordinated expression of phage genes belonging to different temporal classes.

## 1. Introduction

During bacteriophage infection, the expression of viral genes is tightly regulated to ensure that viral proteins are produced at the time when they are needed. Much of this regulation occurs at the level of transcription of viral genes. For most phages, three temporal classes of genes can be defined: early, middle, and late, according to the time after the infection when their transcription begins. Early genes products typically orchestrate the take-over of the host by the virus, middle genes typically encode enzymes involved in phage genome replication, and late genes encode virion and host lysis proteins.

While the tripartite strategy of phage gene expression is broadly applicable to most phages, it is accomplished in very different ways. Some phages rely on the host RNAP to transcribe their genes. In this case, phage-encoded proteins direct the host RNAP to appropriate promoters throughout the infection [1,2] and/or phage-encoded antitermination factors allow host RNAP to access specific groups of phage genes located downstream of terminators [3–5]. Prototypical examples of such strategies are bacteriophages T4 and λ, correspondingly. Some phages encode their own RNAPs. These enzymes are frequently the products of early genes transcribed by host RNAP from strong promoters and are responsible for middle and/or late viral transcription [2,6]. A phage-encoded inhibitor of the host enzyme commonly accomplishes a switch from host to viral RNAP transcription. A prototypical example of such strategy is bacteriophage T7 [6]. Another example is bacteriophage N4, which uses two viral single-subunit polymerases and a host polymerase for its genome transcription. One RNAP is injected inside the cell with phage DNA and transcribes early genes [7,8]. Another RNAP is encoded by early genes and transcribes middle genes [9,10]. Late N4 gene transcription is performed by host RNAP [11].

Transcription of jumbo bacteriophages similar to *P. aeruginosa* phage phiKZ is completely independent of host RNAP [12–14]. These phages encode two multisubunit RNAPs [12–15]. One RNAP is located in the virion (vRNAP) and must be injected into the host cell along with phage DNA. It transcribes early phage genes. The second RNAP, composed of early genes products, is not present in virions (nvRNAP) and recognizes promoters of late viral genes [15,16]. Some phiKZ genes belong to the middle transcription class. It is unknown which of the two viral enzymes is responsible for their transcription. Also, it is unknown how transcription between different classes of jumbo phage genes is switched.

Immediately after the phiKZ DNA injection, a round compartment (RC) appears in the cell [17]. Presumably, RC contains phage DNA and co-injected virion proteins. Later in infection, a large nucleus-like structure is formed in the middle of the cell [17,18]. The phage nucleus (also referred to as pseudo-nucleus) consists of a shell made of phage-encoded protein PhuN [19–21] with replicating phage DNA is located inside [17,18,20]. For phiKZ-related phage 201phi2-1, it was shown that some phage proteins required for transcription and replication, including two subunits of its nvRNAP, are also localized inside the nucleus [22].

In this work, we studied the dynamics and localization of phiKZ RNAPs in infected cells. Our results show that in the very beginning of infection vRNAP co-localizes with injected phage DNA. The vRNAP accumulated throughout the infection is localized in infected cell cytoplasm and is excluded from the nucleus. In contrast, nvRNAP is concentrated inside the nucleus. We propose that such compartmentalization ensures coordinated expression of phage genes during the infection.

## 2. Materials and Methods

### 2.1. Bacteriophage, bacterial strain and growth conditions

The phiKZ phage lysate was prepared as described previously [17]. Phage was further purified using PEG8000 purification protocol [21].

The *Pseudomonas aeruginosa* strain PAO1 culture was grown in a LB medium at 37°C. To prepare infected cells samples, an overnight culture was diluted 1:100 in 1 liter of fresh LB medium. Growth was continued until OD600 reached of 0.6 and phiKZ was added at a multiplicity of infection of 10. Aliquots were withdrawn at selected time points and infection was terminated by the addition of 100 μg/ml chloramphenicol and rapid cooling in an ice water bath. Cells were harvested by centrifugation (5,000 g for 5 minutes), flash-frozen, and stored at -20 °C until use. The infection efficiency was checked by determining the number of colony forming units just before the infection and in infected cultures 5 minutes post-infection. Only cultures that contained less than 20% of surviving cells were used for further processing. For fluorescent microscopy experiments, PAO1 cells were transformed with plasmid expressing fusion protein genes as described carbenicillin [24] and selected on LB agar plates containing 100 μg/ml carbenicillin [24]. A single transformed colony was grown overnight at 37 °C in liquid LB medium supplemented with 100 μg/ml carbenicillin. The culture was diluted 1:100 in fresh LB medium containing carbenicillin, grown at 37 °C until OD600 reached 0.2 and induced with arabinose (final concntrations of 0.1 or 0.2%). Growh was continued at 30 °C until OD600 reached 0.6. At this point, culture aliqouts were used for fluorescent microscopy experiments or for phage infection to obtain containing fusion proteins fluorescent phage progeny.

### 2.2. Cloning

All oligonucleotides use for cloning and/or PCR are listed in the Supplementary Table S1. Gene *180* was amplified from DNA purified from infected cell cultures collected 30 minutes post-infection and cloned into the pET28a plasmid polinker between the NdeI and BamHI sites. The gene of the mCherry fluorescent protein was inserted into the pHERD20T plasmid according to the manufacturer’s protocol of NEBuilder® HiFi DNA Assembly kit (New England Biolabs, UK). The resulting plasmid was named pHERDmCh. To create pHERDgp55-mCh, PhiKZ gene *55* was amplified from the pET28aHis-gp55 plasmid encoding histidine-tagged Gp55 [25] and inserted into pHERDmCh using the Gibson cloning method. The optimized for expression in PAO1 cell gene gp180 was synthesized by IDT and inserted into pHERDmCh using NEBuilder® HiFi DNA Assembly kit (New England Biolabs, UK).

### 2.3 Proteins expression and purification

Rosetta(DE3) *E. coli* cells were transformed with pET28aHis-gp55 or pET28aHis-gp180. Expression was induced by the addition of 1 mM IPTG to cultures grown to OD600 = 0.5-0.7 and further growth at 15°C (Gp55) or 22°C (Gp180) for 5 h. 2 g of wet biomass was disrupted by sonication in 20 ml of buffer A (40 mM Tris-HCl, pH 8.0, 10% glycerol, 500 mM NaCl, 1 mM DTT, 5 mM imidazole) followed by centrifugation at 11,000 g for 30 min. Clarified lysate was loaded onto a HisTrap HP 5 ml (GE Healthcare Life Sciences, USA) column equilibrated with buffer A and washed by the same buffer. The recombinant proteins were eluted with buffer B (buffer A with containing 250 mM imidazole). The eluted fractions were concentrated on Amicon Ultra-4 Centrifugal Filter Unit with Ultracel-10 membrane (EMD Millipore, Merck, USA) and further purified by size exclusion chromatography using a Superdex 200 Increase 10*/*300 (GE Healthcare Life Sciences, USA) in TGED buffer (20 mM Tris–HCl pH 8.0, 5% glycerol, 0.5 mM EDTA, 1 mM DTT) with 200 mM NaCl. Protein concentration was determined using the Bradford assay.

### 2.4 Western blotting

Purified His-Gp55 and His-Gp180 samples were used for rat immunization. Immune antisera were used as primary antibodies at 1:1000 dilution. Peroxidase Goat Anti-Rat IgG (Jackson ImmunoResearch, USA) was used as secondary antibody. Peroxidase activity was detected with SuperSignal™ West Pico Chemiluminescent Substrate (Thermo Scientific, USA). For mCherry and mCherry-fused proteins detection, Anti-tRFP primary antibodies (Evrogen, Russia) were used with Goat anti-rabbit IgG-peroxidase conjugate (Sigma Aldrich, USA) for secondary detection. Western blotting was performed using Amersham Protran 0.45 NC nitrocellulose membranes (GE LifeScience, USA).

To test antisera cross-reactivity, Western blotting was performed with PEG-purified phiKZ virions (10^9^ plaque forming units) and 0.5 μg purified native nvRNAP (Figure S1a). To analyze the vRNAP and nvRNAP subunits synthesis dynamics, three independent cell infection experiments were performed. For each experiment, 100 μl of cell culture before the infection, and 5, 10, 15, 20, 30, and 40 minutes post-infection was used. After probing with the anti-Gp180 sera, nitrocellulose membranes were stripped according to the manufacturer’s protocol and re-probed with the Gp55 antisera. To estimate Gp180 amounts in virions and in 30-minute infected cells, dilutions of purified His-Gp180 of known concentration were used as calibrants. Blots were scanned using the ChemidocTM XRS+ system (Biorad, USA), and individual band densities were measured and compared using Quantity One 1-D analysis software (Biorad, USA). The resulting values were normalized separately for each set of time points.

### 2.5 Co-immunoprecipitation

All procedures were performed at 4°C. Cell pellets from 200 ml of infected cultures were resuspended in 1 ml of buffer C (40 mM Tris-HCl pH 8.0, 150 mM NaCl, 0.5 mM EDTA, Glycerol 10%, 1 mM PMSF) and disrupted by sonication. Crude extracts were clarified by centrifugation at 10,000 g at 4°C. 10 μl of Anti-mCherry ABs solution (Abcam) was added and the lysate was incubated for 1 hour on a rotary shaker. The mixture was transferred to Micro Bio-spin columns (Bio-Rad, USA) containing 250 μl of A-sepharose (GE LifeScience, USA) equilibrated with buffer C. The column was incubated on a rotary shaker for 1 hour. Next, A-sepharose was washed with 10 ml of buffer C and 3 ml of PBS buffer. Proteins were eluted with 250 μl of elution buffer (0.2% SDS, 0.1% Tween-20). The eluted proteins were precipitated with trichloroacetic acid. The protein pellet was resuspended in Laemmli’s buffer and resolved by 10 % SDS-PAGE.

### 2.6 Mass-spectrometry analysis

Protein bands of interest were manually excised from the Coomassie-stained SDS– polyacrylamide gels. Individual slices were prepared for mass spectrometry by *in situ* trypsin digestion at 37°C for 4 h as previously described [15].

### 2.7 Fluorescence microscopy

Cells were withdrawn from infected cultures and immediately placed on microscopic slides, prerared as described in [26]. In the case of fusion proteins expression, agarose pads contained 0.2% (mCh-Gp180) or 0.1% (Gp55-mCh) arabinose. When cells were absorbed, the agarose pad was sealed by a cover glass and VALAP (lanolin, paraffin, and petrolatum 1:1:1). Fluorescence microscopy was performed using a Nikon Eclipse Ti-E inverted microscope equipped with a custom incubation system. During microscopy, the incubation temperatures were 30 °C for Gp55-mCh expressing cells and 37 °C for other cells. Fluorescence signals in DAPI and mCherry fluorescence channels were detected using Semrock filter sets DAPI-50LP-A and TxRed-4040C, respectively. Imaging was performed using Micro-Manager [27] with a custom script. Images were taken every 10 min for 180 min. Image analyses were performed using the ImageJ software.

The fluorescence intensity corresponding to a single mCherry molecule was calculated as described in *[26]*. First, the fluorescence of pure mCherry protein solution with known concentration was measured using spectrofluorometer and intensity corresponding to single mCherry molecule was calculated. Next, fluorescence of an aliquot of a culture containing known number (CFUs) of *E. coli* cells producing mCherry was measured using spectrofluorometer and the average number of fluorescent protein molecules per cell was determined. Individual cells from the same culture were next observed under the fluorescent microscope. Average fluorescence intensity of single cells was divided by the estimated average number of mCherry molecules per cell. In this way, the fluorescence intensity of a single mCherry molecule was obtained under the imaging conditions used. Since for various experiments the imaging parameters such as exposure, the presence of filters, etc. differed, calibration coefficients were obtained for each imaging condition. These coefficients were calculated by determining the average fluorescence intensity of at least 400 mCherry producing *E. coli* cells at a given imaging condition.

## 3. Results

### 3.1. Dynamics of phiKZ RNAPs accumulation during infection

To analyze the dynamics of phiKZ RNAPs accumulation during infection, we used sera containing polyclonal antibodies (ABs) specific to the Gp180 subunit of vRNAP and Gp55 of nvRNAP (Figure S1a). Both subunits contain the universal Mg^2+^-binding domain DFDGD, which is essential for RNAP activity *[28, 29]* and thus serves as a good proxy for measuring the amounts of active viral RNAPs. The number of vRNAP molecules in the virion was estimated to be 10 (±3) molecules based on the results of semiquantitative analysis using recombinant Gp180 with known concentration for calibration (Figure S1b). Gp180 was detected inside the cells 5 minutes post-infection and its amount steadily increased at later time points up to 30 minutes post-infection and remained constant (Figure 1a, note that infected cells start to lyse after 50 minutes post-infection). At 30 minutes post-infection, the number of Gp180 molecules was estimated as ∼3000 per cell (Figure S1c). Given the reported phiKZ burst size of 100 progeny phage particles per cell *[30]*, this result provides an independent upper estimate of ∼30 vRNAP molecules per virion (note that not all vRNAP molecules may be packaged).

**Figure 1:**
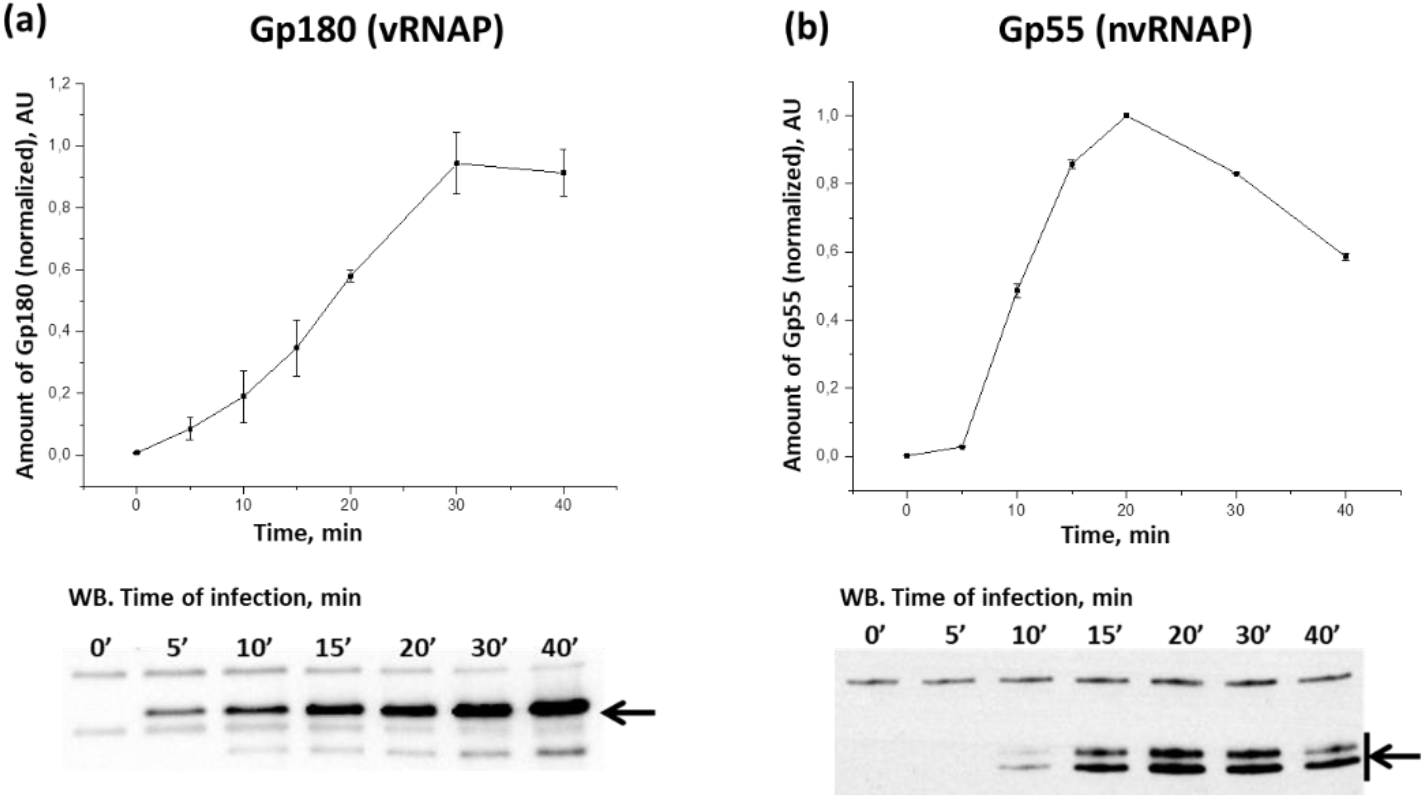
Accumulation of PhiKZ Gp180 and Gp55 in the course of the infection. Changes in RNAP subunits amounts (determined by the intensity of bands on Western blots such as the ones shown below) are presented. For each time point, means from three independent measurements are plotted, error bars represent standard deviations. Black arrows indicate the positions of Gp180 (a) and Gp55 (b) bands; Gp55 migrates as two bands.

Gp55 (and, presumably, nvRNAP) appeared 10 min post-infection. Its amount reached a maximum 20 minutes post-infection and slightly decreased at later times (Figure 1b).

### 3.2. Determination of localization of phage RNAPs during infection using fluorescence microscopy

To monitor the intracellular localization of phage RNAPs within infected cells, we constructed expression plasmids carrying genes encoding phage RNAPs subunits fused with the mCherry fluorescence protein (pHERD20TmCherry-gp180 and pHERDStr1gp55-mCherry, see Materials and Methods). Using Western blots, we showed that (1) both fusion proteins were produced inside *P. aeruginosa* cells transformed with either of these plasmids, and (2) there was no degradation of fused proteins in either uninfected or infected cells (Figure S2). Live fluorescent microscopy revealed that fusion proteins and the control mCherry protein were diffusely distributed in uninfected cells. In infected cells, mCherry and mCherry-Gp180 were localized in the cytoplasm and were absent from the phage nucleus. In contrast, mCherry-Gp55 accumulated inside the nucleus (Figure 2).

**Figure 2.**
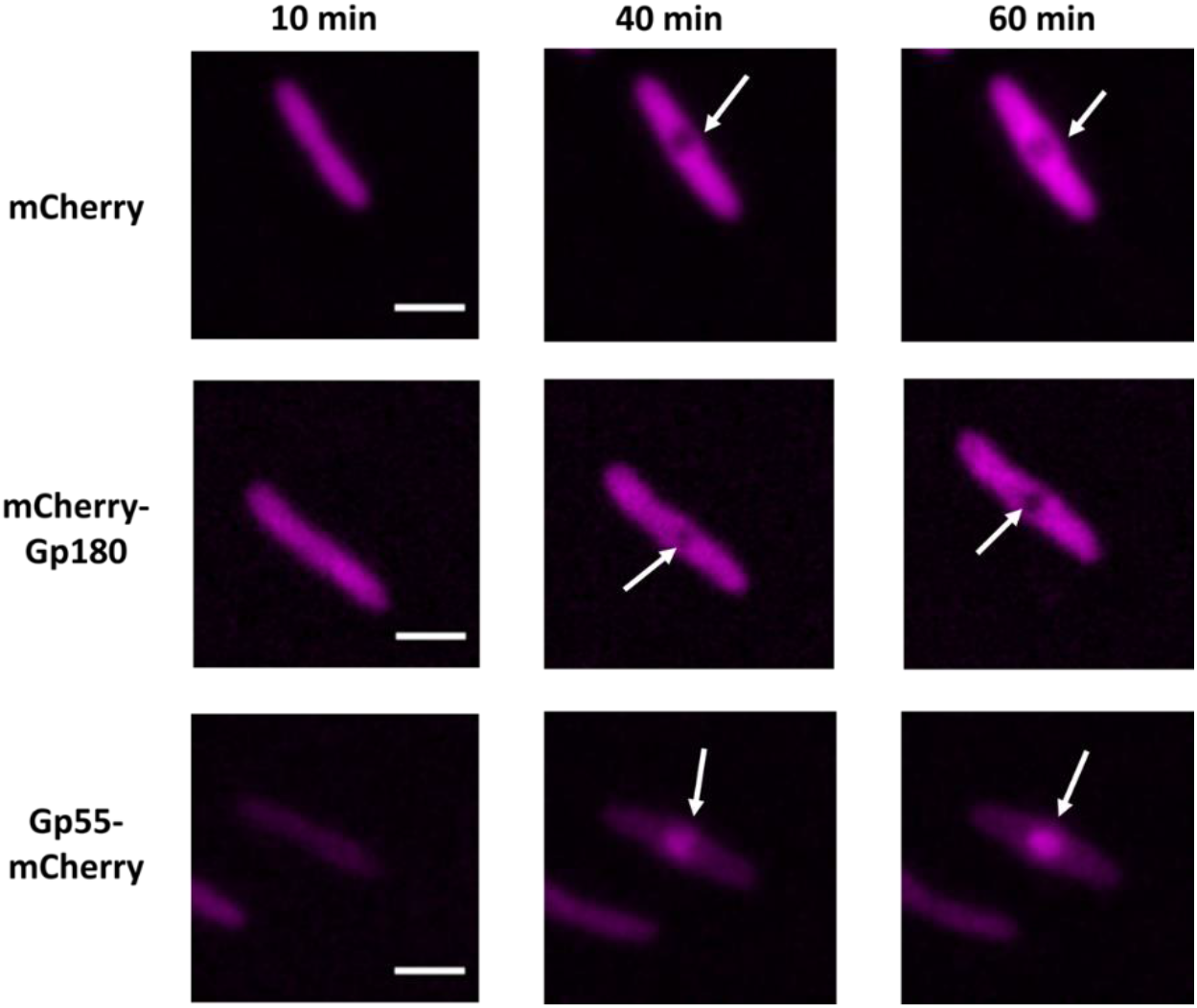
Localization of mCherry fluorescent protein, and Gp180 and Gp55 mCherry fusions during the PhiKZ infection. White arrows point to the phage nucleus. Time indicates minutes post-infection. All scale bars are 2 μm.

Since mCherry-Gp180 localized just like the mCherry control throughout the infection, it was necessary to show that its distribution reflects localization of vRNAP. We conducted immunoprecipitation with mCherry antibodies using extracts of infected and uninfected cells. In the infected cell sample, we detected vRNAP subunits Gp178, Gp80, and Gp149 along with mCherry-Gp180 (Figure S3a), indicating that the fused subunit indeed is assembled into the vRNAP complex. We also collected phage progeny formed after the infection of mCherry-Gp180 producing cells and analyzed the protein content of progeny virions by Western blotting (Figure S3b) and fluorescent microscopy (Figure 3a). As a control, we used phages released from cells expressing the mCherry protein. The results showed that mCherry-Gp180 was incorporated into virions. Earlier, the coefficients for conversion of fluorescence intensities of individual cells to the number of mCherry fluorescent protein molecules were determined *[26]*. We used this information and fluorescent data for 15 virions to estimate the number of mCherry-Gp180 molecules per virion. Our measurements provided an estimate of 7±4 molecules per particle (Figure S4), which *i*) is consistent with the number obtained using Western blotting (above) and *ii*) should be considered as a lower boundary since non-fluorescent vRNAP with Gp180 encoded by the phage may also be present in the virion.

**Figure 3.**
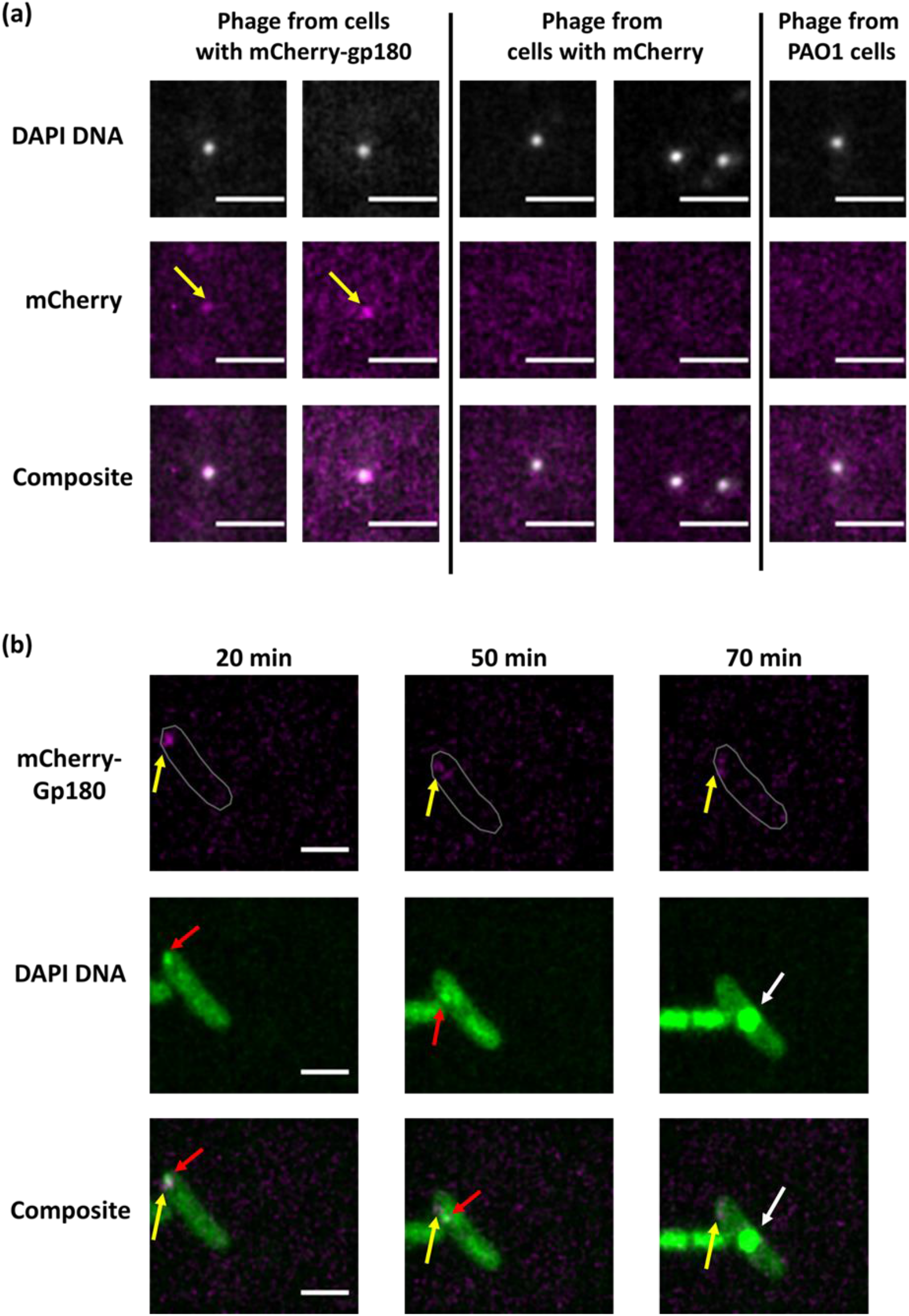
Localization of PhiKZ vRNAP in infected cells. (a). Co-localization of DAPI stained DNA and the mCherry signals in PhiKZ phage particles in the lysate of mCherry-gp180 containing cells revealed by fluorescent microscopy. The scale bars are 1 μm. Phage particles from the lysate of mCherry-containing and native PAO1 strain cells are shown as controls. (b). Images of PAO1 cells infected with mCherry-Gp180 containing phiKZ phage. Yellow arrows point to mCherry-Gp180, red arrows point to the area of concentrated DNA (presumably RC), and white arrows – to the phage nucleus. Time indicates minutes post-infection. Cell contour is delineated (gray). The scale bars are 2 μm.

Next, we used phages containing mCherry-Gp180 to infect *P. aeruginosa* cells and followed the infection by live fluorescence microscopy. As is shown in Figure 3b, the mCherry signal appeared in infected cells early in infection and co-localized with a bright DAPI-staining area (presumably corresponding to phage DNA located inside RC). Notably, we detected DAPI-dot near the cell pole at the beginning of infection. Later in infection, when the phage nucleus formed in the middle of the cell, no mCherry-signal was detected inside of it. We conclude injected vRNAP co-localizes with injected phage DNA at the earliest stages of infection and that both injected and newly synthesized vRNAP are excluded from the phage nucleus.

## 4. Discussion

In this work, we show using several independent methods that the number of vRNAP molecules per phiKZ virion is between 10 and 30, which is comparable to the number of early phage promoters [12,31]. We also show that phiKZ vRNAP is injected into infected cells and co-localizes with the phage genome in the round compartment (RC) at the earliest stages of infection. The The RC is likely separated from the cytoplasm DNA [17], which must help compartmentalize early transcription and make it more efficient. The products of early genes are needed for phage nucleus formation and also include the nvRNAP subunits. Once the injected PhiKZ genome finds itself in the nucleus it begins to be replicated [17]. Transcription of phage genomes enclosed in the nucleus is performed by the nvRNAP, which is localized in this compartment. In contrast, the injected vRNAP remains excluded from the nucleus, providing an elegant and simple mechanism of a switch from early transcription to middle and late phiKZ transcription initiates from promoters with different consensus sequences [12]. Given that vRNAP is encoded by middle genes [12], and that at least *in vitro*, the phiKZ nvRNAP transcribes from late promoters only [15,25], it remains to be determined which enzyme, likely modified by additional DNA-or RNAP-binding factors, is responsible for middle transcription, which presumably happens when the nucleus is being formed.

Additional vRNAP molecules are assembled throughout the infection upon translation in the cytoplasm. The newly synthesized vRNAP remains in the cytoplasm, solving yet another problem of preventing early transcription from accumulating phage genomes. Eventually, these vRNAPs must be packaged into progeny phage particle heads that are assembled near the cell wall and then transferred to the cytoplasmic face of the nucleus to be packed with phage DNA [32]. Gp219, a protein of unknown function that we found to be associated with vRNAP in infected cells (Supplementary Figure S3), is absent from the phiKZ virions [33,34] but may play a role in vRNAP packaging.

Overall, our results provide evidence that the jumbo phages nucleus allows, in addition to avoiding host defense mechanisms [35], to spatially separate transcription by two phage-encoded RNA polymerases, ensuring temporal coordination of phage transcription. Some jumbo phages do not encode PhuN homologs [29] and develop without the formation of a nucleus compartment in infected host cells. Invariably, the DNA of such phages contains uracyl instead of thymine, which presumably allows to avoid host defenses [13,36]. Uracyl-containing jumbo phages also encode two multisubunit RNAPs that are similar to nucleus-forming phages transcription enzymes [13,29]. It should be highly interesting to determine how coordinated temporal transcription of these uracyl-containing genomes is achieved in the absence of separate compartments.

## Supplementary Materials

The following supporting information can be downloaded at: www.mdpi.com/xxx/s1, Figure S1: (a) The analysis of cross-reactivity between serums (Abs) to vRNAP (Gp180) and nvRNAP (Gp55). The examples of western blots for semiquantitative analysis of the vRNAP amount in the single phage particle (b) and on the 30th minute of infection (c).; Figure S2: The result of analysis of recombinant phage RNAPs subunits Gp55-mCh (a) and mCh-gp180 (b) expression and stability during infection.; Figure S3: Confirmation of the inclusion of mCh-Gp180 in new virions as part of vRNAP; Table S1: The list of oligonucleotides, which were used for cloning.

## Author Contributions

Conceptualization, K.S. and M.V.Y.; investigation, D.A., V.V.B., E.Zh., M.S., I.E.V., T.A., I.K., N.M., A.A. and M.K.; writing—original draft preparation, M.V.Y and V.V.B.; writing—review and editing, M.V.Y., K.S. and M.Kh.; visualization, D.A., N.M. and M.S.; funding acquisition, M.V.Y. and K.S. All authors have read and agreed to the published version of the manuscript.

## Acknowledgments

This research was carried out using scientific equipment of the Center of Shared Usage «The analytical center of nano- and biotechnologies of SPbPU.

