## Supplementary material for "The Dynamics of Synthesis and Localization of Jumbo Phage RNA Polymerases inside Infected Cells"

**Supplementary materials**

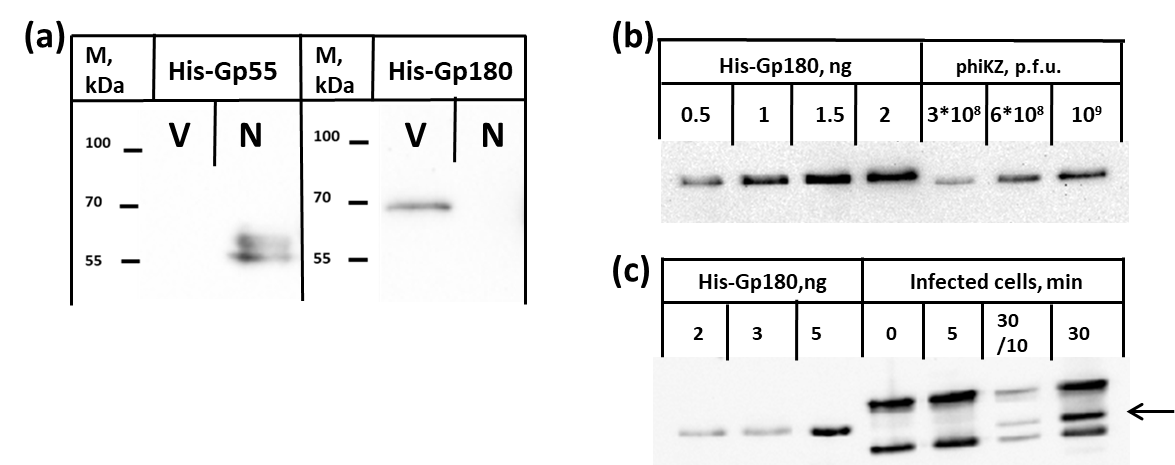

**Figure S1**: (a) Antibodies (ABs) raised against vRNAP His-Gp180 and nvRNAP His-Gp55 are specific and not cross-reactive with PhiKZ RNAPs. Purified phiKZ virions were used as a sample containing vRNAP only (labeled “V”). Purified nvRNAP isolated from PhiKZ-infected cells ("N") was used as sample containing nvRNAP only. The samples were separated by SDS PAGE and probed with antisera raised against vRNAP His-Gp180 and nvRNAP His-Gp55. (b) A representative Western blot used to semi-quantitatively estimate vRNAP amounts in virions. The right three lanes show signals obtained in samples containing indicated number of infectious PhiKZ virions (plaque forming units). Four left lanes show signals from indicated amounts of pure recombinant hexahistidine-tagged Gp180. (c) A representative Western blot used to estimate vRNAP amounts inside infected cells. Three lanes on the left show signals from indicated amounts of pure recombinant hexahistidine-tagged Gp180. Four lanes on the right show signals obtained with extracts of cells infected at a multiplicity of 2 collected at indicated times post-infection. Cross-reacting bands above and below the Gp180 bands are present in both uninfected and infected cells and are coming from unidentified cell proteins. The two rightmost lanes show Gp180 signals from the standard amount of 30 minute post-infection extract used in other lanes (rightmost lane) and from the same volume of 10-fold diluted extract (second from right lane).

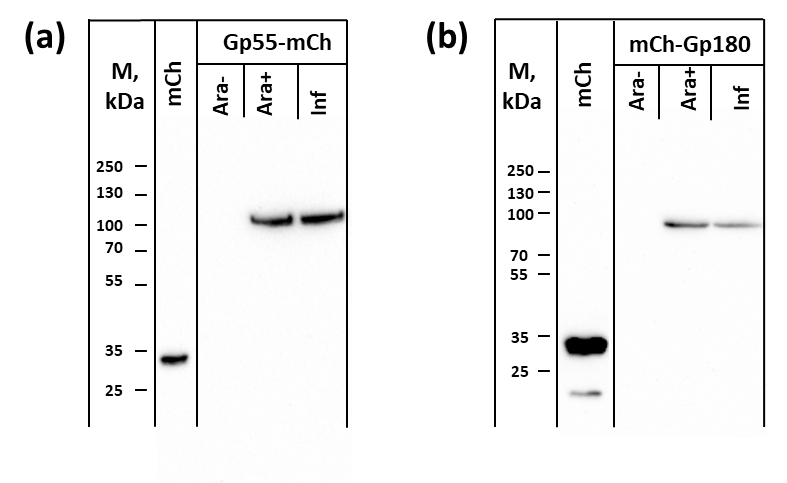

**Figure S2**. The mCherry fusions with nvRNAP Gp55 (a) and vRNAP Gp180 (b) are stable throughout the PhiKZ infection. “mCh” – recombinant mCherry protein used as a control. “Ara-“ - samples of cells containing an expression plasmid with indicated fused protein gene grown in the absence of arabinose inducer; “Ara+” - induced samples immediately prior to infection with PhiKZ; “Inf” - induced samples infected with PhiKZ at high MOI and analyzed 30 minutes post-infection.

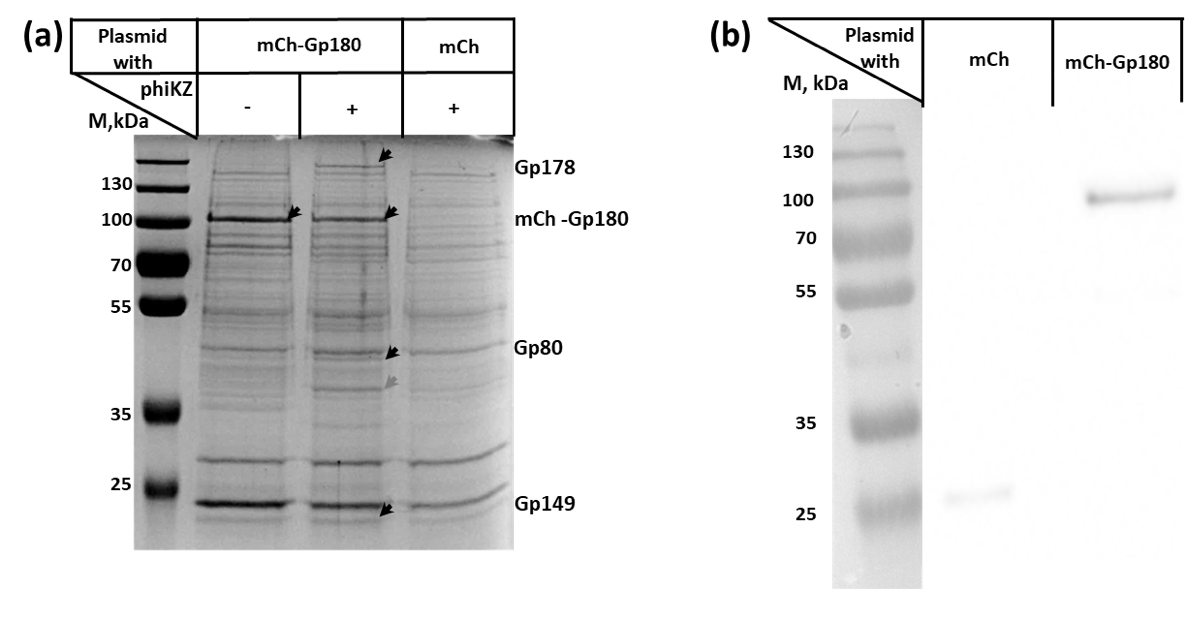

**Figure S3**: The mCh-Gp180 fusion is incorporated in vRNAP and progeny phage virions. (a) The result of co-immunoprecipitation with anti-mCherry antibodies from non-infected cells expressing mCherry-Gp180 fusion or phiKZ-infected cells expressing the mCherry-Gp180 fusion or mCherry alone (“mCh”) collected 30 min post-infection. Protein bands enriched in infected cells expressing mCh-Gp180 with or without infection were analyzed by mass-spectrometry. Black arrows point to protein bands identified on the right. The grey arrow indicates Gp219, an early PhiKZ protein of unknown function. (b) Western-blot analysis of phiKZ progeny formed after infection of expressing mCherry or mCh-Gp180 fusion. Material corresponding to 10^9^ plaque forming units was loaded on each lane. The weak signal from the mCherry protein in the control sample is attributed to either spurious incorporation of mCherry into virions or its binding to virion’s surface.

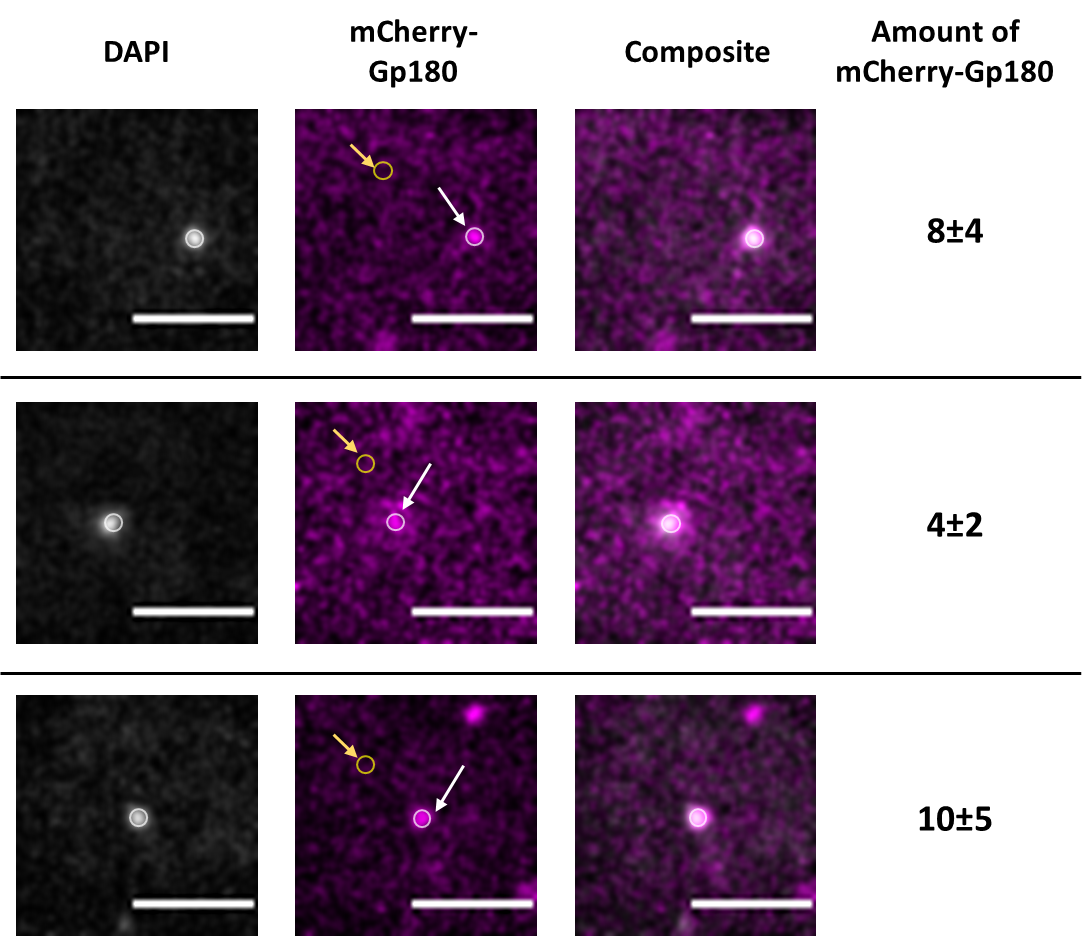

**Figure S4:** Representative examples of individual phage particles with the mCherry-Gp180 fusion protein inside. Phage particles formed after infection of mCherry-Gp180 overproducing cells were first imaged in the DAPI fluorescence channel, and 60x60 pixels squares containing individual phage particles were selected and duplicated. An area matching with a virion in the DAPI fluorescence channel (white circle and white arrows) was selected by the oval selection tool and fluorescence intensity in the red channel was measured. To measure background intensity, fluorescence was measured in the same-sized area that had no phage particles. Positions of background measurements are marked with yellow circles with yellow arrows in red fluorescence channel images (middle). Background intensity was subtracted from mCherry-Gp180 fluorescence intensity. Estimated numbers of mCherry-Gp180 molecules for each virion are presented for each of the three representative virions shown. The scale bar is 2 μm.

**Table S1. The list of oligonucleotides used for cloning.**

| **Oligonucleotide name** | **Sequence** |
| --- | --- |
| pHERD20TG-F | GGGTATGTATATCTCCTTCTTAAAG |
| pHERD20TG-R | CACGTGCCATGGGATCTGATAAG |
| pHERD20TGp180Opt-GR | ATGAAGAAAGCGCTGGTGCC |
| mCherryG-R | CTTATCAGATCCCATGGCACGTGCTTGTACAGCTCGTCCATGC |
| mCherryG-F | CTTTAAGAAGGAGATATACATACCCATGGTGAGCAAGGGCGAGG |
| KZ180-F | AGGCATATGAAGAAGGCATTAGTTCC |
| KZ180-R | CTGGATCCTTAATCCCTTTGTTTGG |
| pHerdt-mCh-GR1 | TCGGTACCCGGGGATCCTC |
| KZGp55F-G | AAGGAGATATACATACCCATGGGTCTTTATGCCAAAG |
| KZGp55-RminuSt-G | GAGGATCCCCGGGTACCGATTTTGCTTTTAGCCATTCTTC |
